# GemSpot: A Pipeline for Robust Modeling of Ligands into CryoEM Maps

**DOI:** 10.1101/750778

**Authors:** Michael J. Robertson, Gydo C. P. van Zundert, Kenneth Borrelli, Georgios Skiniotis

## Abstract

Producing an accurate atomic model of biomolecule-ligand interactions from maps generated by cryo-electron microscopy often presents challenges inherent to the methodology and the dynamic nature of ligand binding. Here we have developed GemSpot, a pipeline of computational chemistry methods that take into account EM map potentials, quantum mechanics energy calculations, and water molecule site prediction to generate candidate poses and provide a measure of the degree of confidence. The pipeline is validated through several published cryoEM structures of complexes in different resolution ranges and various types of ligands. In all cases, at least one identified pose produced both excellent interactions with the target and agreement with the map. GemSpot will be valuable for the robust identification of ligand poses and drug discovery efforts through cryoEM.

## Introduction

Electron Cryo-Microscopy (cryoEM) has emerged as a major methodology for high-resolution structure determination of macromolecules and their complexes. The number of deposited cryoEM structures in the PDB^1^ with resolution better than 4 Å has increased from 48 in 2015 to almost 1,500 to date. These structures include many macromolecular complexes and membrane proteins that have generally proven very challenging or intractable for traditional structural techniques, particularly X-ray crystallography. For example, GPCRs and ion channels are very important classes of drug targets representing 33% and 18% of FDA approved pharmaceuticals^2^, respectively, where structural studies have been historically limited by difficulties associated with their crystallization, although recent advances have been made^3^. CryoEM, on the other hand, offers increasingly robust workflows for the structural determination of these types of macromolecules^4^. As a result, cryoEM is opening unprecedented opportunities for structure-based drug discovery on a large variety of targets that were up to recently intractable.

Structure-based drug discovery is a rational drug design approach that takes into account the three-dimensional structure of the biomolecular target^5^. The unliganded structure can be employed for large-scale virtual screening to get initial hit compounds with the desired biochemical activity^6^. With lead compound(s) in hand, an experimental structure of the liganded complex is necessary to verify the exact binding mode, and often assists in identifying modifications to improve potency. The correct pose is particularly important for methods such as free energy perturbation calculations, where a compound is alchemically mutated to an analogue over the course of a molecular simulation and the relative free energy of binding is calculated, as was recently successfully shown on a cryoEM derived structure for human ATP-citrate lyase^7^. Furthermore, a sufficiently high-resolution (usually <2.5 Å) structure will allow for the identification of bound water molecules, which can play crucial roles in drug design^8^. For example, development of successful HIV protease inhibitors often involves the replacement of a key structural water^9^. Given the recent remarkable progress of cryoEM, the methodology will become an invaluable tool for drug discovery efforts, especially for challenging macromolecular complexes. Underlined by continuous advancements in sample preparation^10^, automated data collection^11^, and improved availability of microscopes capable of achieving high resolution, cryoEM will inevitably be employed in the lead optimization phase to obtain structures of intermediate compounds bound to their targets.

Decades of crystallography have led to robust methods of modeling and validating protein-ligand crystal structures^12^. While both X-ray crystallography and single-particle cryoEM are in principle scattering techniques based on the interaction of radiation with a biological specimen, there are key differences that complicate modeling in cryoEM maps and prevent the usage of the metrics developed for crystal structures. In crystallography, the phase information of the scattered radiation that is measured is lost and needs to be recovered with either additional experimental information (*e.g.,* Multi-wavelength anomalous dispersion (MAD), isomorphous displacement) or comparison to known structures (molecular replacement)^13^. The initial phase values are then improved during model building by comparing calculated scattering from the current model to the experimental scattering. Thus, X-ray crystallography structure determination involves a continuous cross-talk between model and experimental data with simultaneous feedback on the quality of the model. By contrast, in cryoEM the phases are readily available as they are embedded in the specimen images, which are directly used for the calculation of 3D maps. Once a final threedimensional map has been determined from thousands of experimental projections, the model is built into the map with no further feedback from the raw EM data. Furthermore, the maps obtained in crystallography correspond to the electron density, while in cryoEM they represent the coulombic potential of the molecule under investigation. Thus, using the tools developed for crystallography directly for cryoEM structure modeling can be inherently problematic.

While the number of cryoEM maps of macromolecular complexes determined to date is relatively low, the existing structures suggest that there are some fundamental challenges associated with modeling protein-ligand complexes. Even with very high-resolution data for a biomolecule, the resolution of the map for a bound ligand is often significantly lower than its surrounding environment^14^. Given that cryoEM structures derive from flash-frozen macromolecules in aqueous solution, it is perhaps not surprising to observe additional mobility for some ligands within protein active sites. In addition, cryoEM reconstructions are vulnerable to spurious map features, currently evident with different software yielding noticeably different maps from the same dataset. This characteristic may arise from inaccuracies in image defocus estimation and correction of the contrast transfer function at high resolution, as well as variability in masking and weighting schemes employed in different software platforms for processing cryoEM data. Notably, in some cases, even different settings with the same software will yield map deviations that may have significant effects in ligand modeling. This problem is compounded by the fact that ligands lack the structural constraints adopted by proteins, *e.g*., secondary structure constraints that facilitate more robust modeling. Such caveats present the modeler with the challenge of identifying the bound pose of a ligand within a relatively high-resolution cryoEM map, resulting in often incorrect ligand poses and interpretations with significant implications for molecular mechanism and drug discovery efforts.

Parallel to developments in cryoEM, computational chemistry methods for modeling protein-ligand complexes have improved significantly over time. Computational force fields have been successfully used for decades to describe the energy and forces of various conformations of proteins^15^. These force fields have been expanded to accurately describe the energy and force of a large variety of ligands, and can easily be expanded by users to cover ligands of interest or even be automatically extended to cover ligands outside of those used in the initial parameterization^16,17^. Such force field parameters have been used for a variety of applications including dynamics^18^ and for enumerating the conformations of proteins and ligands that will be accessible in biologically relevant conditions^19^. Molecular docking is an approach that uses force fields, in conjunction with highly optimized sampling and refinement algorithms, to predict protein-ligand binding modes given only the conformation of the protein and the identity of the ligand^20^. This methodology has been extensively applied to both identify ligands that bind to specific proteins with high affinity and to predict their protein-ligand binding conformations^21^. It should be noted however that, in the absence of experimental data, these purely computational methods are often hampered by significant false positive and false negative rates.

For structure-based drug design, significant emphasis has also been put on predicting the location of water molecules, which often coordinate ligand binding in pockets and have profound effects in pharmacological activities. Several approaches for predicting hydration sites, including grid-based approaches like JAWS^22^ and dynamics approaches like WATERMAP^23^, are now capable of predicting the location of bound water molecules. These computational predictions yield impressive agreement with experimentally derived structures and further highlight the role of hydration in lead optimization^24^.

It thus becomes apparent that an array of well-established computational tools can be employed in combination with cryoEM to address the challenge of modeling ligands into cryoEM maps. To this end, we have developed and validated ‘GemSpot’, a pipeline of computational chemistry methods that assists in obtaining the most probable bound pose using a combination of ligand docking coupled with refinement, quantum mechanical (QM) calculations, automatic water placement and additional external information, all while taking into account the experimental cryoEM data. The GemSpot pipeline has been validated against a varied set of 19 structures obtained from cryoEM data ranging from 1.9-4.3 Å resolution, consisting of both protein and RNA (Supplementary Table 1), together with a diverse selection of ligands, including small molecules and peptides (Scheme of all ligands in Supplementary Fig. 1).

## Results

### The GemSpot pipeline

In the first step using GemSpot (see Fig. 1), the ligand is docked with the popular software GLIDE^25^ by employing a novel combination of the traditional GLIDE docking score function and a real space cross-correlation score to the map. This software, called GlideEM, generates several candidate poses for the ligand that are then subjected to real space refinement with PHENIX^26^ including the state-of-the-art OPLS3e / VSGB2.1 force field^16,27^. A combination of real space correlation coefficient and pre-refinement docking scores are used to eliminate any poses that make little chemical sense or fit poorly into the experimental map. Once the top poses are identified, further computational techniques can be used to generate enhanced confidence in the lead candidate pose, when necessary. For high-resolution EM maps, a free energy approach to hydrate the active site using JAWS^22^ can be used to help differentiate potential water molecules from noise in the map and gain insight into ligand interactions. When there are still doubts about the conformation of the molecule, one can leverage quantum chemistry to examine the conformational strain associated with any bound poses, e.g. with GAUSSIAN^28^ or Jaguar^29^. In situations where these computational methods alone may be unable to determine a single pose that unambiguously fits all of the data, it may be necessary to determine which of the top poses are also consistent with data from other experiments. Particularly valuable is comparison to structure-activity relationship (SAR) data, i.e., whether the prospective pose can effectively explain the changes in binding affinity for analogues of that molecule^7^. By combining the resulting data, a high degree of confidence can often be obtained even with a low resolution or problematic density for the ligand.

**Fig 1.**
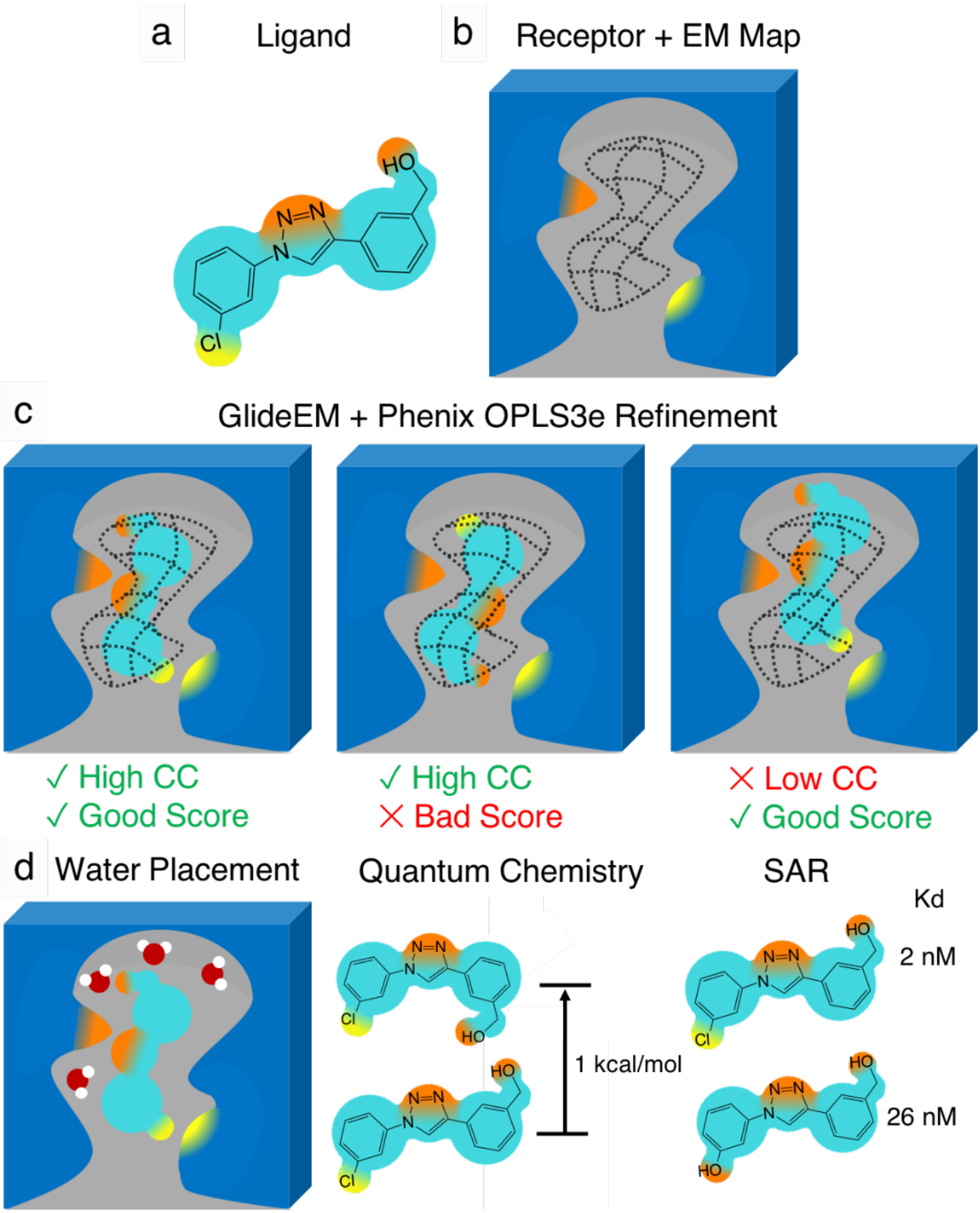
Schematic of the GemSpot pipeline. **a**, Example of a ligand with the potential to make several hydrophobic, hydrogen bonding (orange), and halogen bonding (yellow) interactions. **b**, The ligand’s active site with the ligand EM density map shown in wireframe. **c**, Examples of poses with and without favorable interactions (Good/Bad Score) and/or cross correlation (CC) to the EM map. **d**, Further support for the pose based on water placement, quantum chemistry, and structure-activity relationship data.

### Using GemSpot for beta-galactosidase

The case of beta-galactosidase bound to phenylethyl β-d-thiogalactoside (PETG) provides perhaps the most striking demonstration for this system of modeling protein-ligand complexes using GemSpot. Two published structures, PDB:5A1A^30^ and PDB:6CVM^14^, have been derived from the same data set by using different software packages to determine the three-dimensional maps. A 2.2 Å map associated with the PDB:5A1A structure was obtained using RELION^31^, whereas a 1.9 Å map associated with the PDB:6CVM structure was obtained with CisTEM^32^.

Despite derivation from the same data set, the map densities corresponding to the ligand had different features and, as a result, a significant change to the modeled pose for the ligand was made. Thus, a methodology that could predict the same, correct pose, despite differences in features of ligand densities would be highly desirable. To begin processing ligand modeling in beta-galactosidase, we subjected both protein models with all water and ligand molecules removed (metal ions were retained) to PHENIX real-space refinement prior to docking with GlideEM.

Interestingly, docking to both structures/maps yielded ligand poses with the pyranose ring of PETG in the same orientation as shown in PDB:5A1A, which corresponds to the 2.2 Å map (Fig. 2a,b). The cross-correlation of the top real-space-refined docked poses against the 1.9 Å experimental map is actually higher than the associated PDB:6CVM deposited pose (0.73 compared to 0.71). This further suggests that a pose like the one modeled in PDB:5A1A, corresponding to the 2.2 Å map, was the more probable one. With PDB:6CVM, traditional docking without the inclusion of the EM map also yielded this pose. However, docking against PDB:5A1A without the map yielded predominantly poses with the ligand outside the corresponding EM density (Supplementary Fig. 2). These discrepancies seem to result from subtle differences in the position of the sodium and magnesium ions, which can create steric issues in docking the best pose.

**Fig 2:**
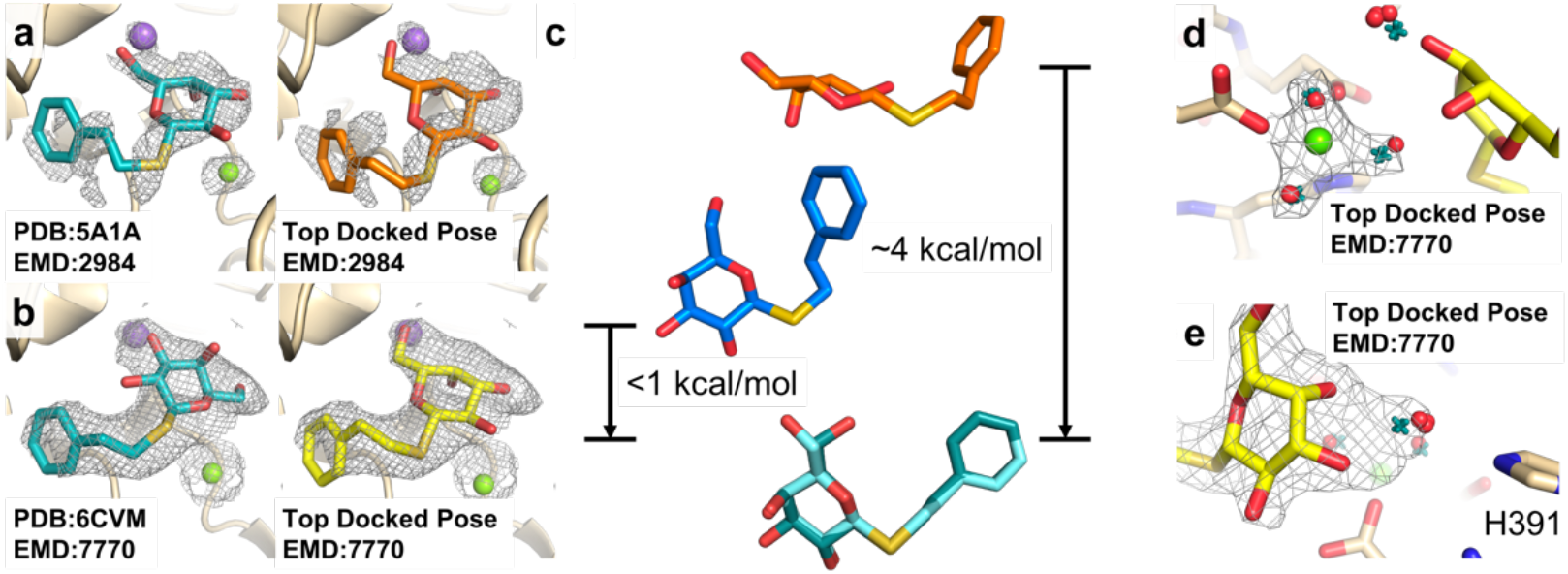
GemSpot results for PETG in beta-galactosidase. **a**, PDB:5A1A with its associated map EMD:2984. The deposited PDB pose is shown in teal, and the best pose obtained from GlideEM is shown in orange. The green and purple spheres correspond to magnesium and sodium ions, respectively. **b**, PDB:6CVM with its associated map EMD:7770. In teal, the deposited PDB pose and in yellow the best pose from GlideEM. **c**, A comparison of the deposited poses from PDB:5A1A (blue) and PDB:6CVM (orange) with the overlaid QM optimized geometries (teal). The energies reported are the difference between that conformation optimized with the O-C-S-C dihedral angle fixed and that conformation optimized without any fixed dihedrals. **d, e**, Results of JAWS calculations performed on the best GlideEM pose for the PDB:6CVM structure. Predicted water sites from triplicate simulations are depicted as red spheres, while real-space refined waters based on those positions are presented as teal crosses, and the map shown is EMD:7770.

While no pose was obtained in any docking test that resembled that deposited in PDB:6CVM, we performed quantum chemical optimizations on the conformation of PETG from PDB:5A1A and PDB:6CVM. Both calculations converged to near-identical conformations (RMSD of 0.18 Å), presented in Fig. 2c. The optimized structure is very similar to that shown in PDB:5A1A, with the exception of the phenyl group extending to a trans conformation. When the optimization began from the conformation shown in PDB:6CVM, we observed a substantial shift in the saccharide ring, with the O-C-S-C dihedral angle shifting from −144 to −71 degrees, the same value as the one in the PDB:5A1A structure. Conformer optimization while maintaining the O-C-S-C dihedral angle fixed at −143 degrees results in a configuration very similar to that of PDB:6CVM. However, this state is roughly 4 kcal/mol higher in energy than the unconstrained state, providing further evidence that the pose modeled in PDB:5A1A is the more physically probable state.

In the next step, we sought to determine water molecule positioning in the structure. For this purpose, we ran JAWS calculations on the pose modeled in Fig. 2b bound to the protein structure from PDB:6CVM, the results of which are presented in Fig. 2d. An octahedral model for magnesium was used to ensure proper coordination of the magnesium ion, as this is the expected coordination of Mg^2+^ and consistent with the features in the density (Fig. 2d). This approach enabled us to predict with high confidence a bound water molecule interacting with PETG and histidine 391. Strikingly, this water molecule resides in part of the density region previously attributed to the ligand alone in PDB:6CVM (Fig. 2d). It thus appears that the observed continuous density may have been the product of close proximity between the ligand and the water molecule, giving rise to uncertainty in modeling that was effectively addressed with the GemSpot workflow. In addition, three sites of hydration were predicted near the sodium ion (Supplementary Fig. 3), with the strongest map features corresponding to the most tightly predicted water site from the triplicate JAWS calculations. By contrast, little consensus was found in the JAWS calculations for the solvent accessible side of the ligand.

In the example of beta-galactosidase, high-resolution crystal structures of the protein in complex with analogous ligands can provide experimental data to assist in determining the correct pose for PETG. Several of these compounds have the same saccharide moiety but differ from PETG in the thiol group. This provides a clear opportunity to examine how this saccharide ring should be correctly modeled in the active site. Comparing the poses from GemSpot, PDB:5A1A, PDB:6CVM, and the 1.6 Å crystal structure of 4-nitrophenyl-beta-D-galactosidase (PNPG) (Supplementary Fig. 4) confirms the results of our modeling, which suggested that a pose much more similar to that of PDB:5A1A corresponds to the correct one. The high-resolution crystal structure also shows the presence of a water molecule, as predicted by our JAWS simulations, which is responsible for the aberrant map feature that led to the modeled ligand pose in PDB:6CVM. Thus, all the evidence points to our best-docked pose as the most probable bound ligand conformation.

### Using GemSpot with ~ 3.0 Å resolution maps

Beyond beta-galactosidase, we examined four other structures with a reported global resolution of 3.0 Å or better. In general, at this resolution it is not uncommon for the human modeler to be able to figure out the correct pose from the map alone. In the deposited structures for cannabinoid receptor 1 (CB1R)^33^, the eukaryotic voltage-gated sodium channel NavPaS^34^, the small subunit of *leishmania* ribosome^35^ and the *leishmania* 20S proteasome^36^ the modeled poses were in excellent agreement with our top calculated pose (Supplementary Fig. 5, Supplementary Table 2). Using JAWS calculations, we were able to not just recapitulate the single water modeled in the NavPaS structure but also suggest two additional hydration sites in the vicinity of the tetrodotoxin ligand that are in fact observable in the EM map (Supplementary Fig. 6).

The small subunit of the *leishmania* ribosome presents an interesting example where docking without the EM map yields predominately poses that do not agree well with the map (Fig. 3), possibly because the large number of positive charges on the paromomycin ligand can match well with the phosphodiester backbone in many locations, with every pose scoring well. However, this problem was resolved when the map densities are included in the docking using GlideEM. While these structures provide a valuable test for the whole pipeline, it is crucial to also examine performance for lower-resolution structures where a human modeler may struggle to model from the EM map alone.

**Fig 3:**
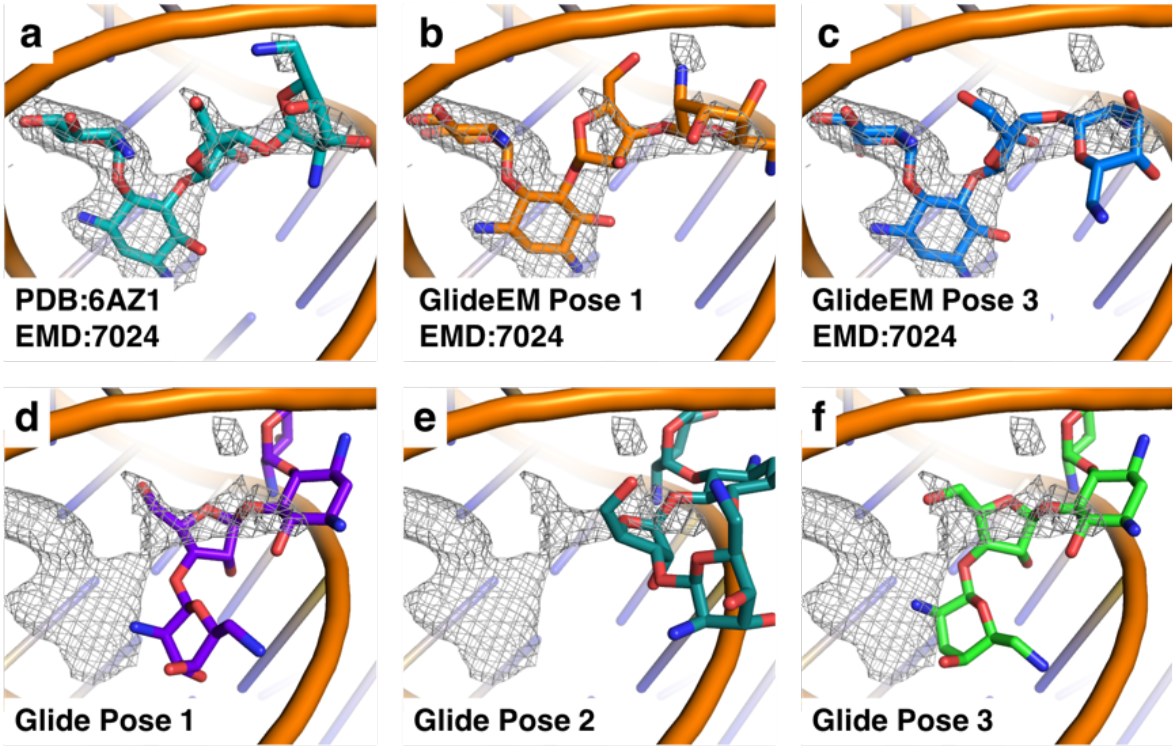
Comparison of paromomycin poses docked with and without the EM map. **a**, PDB: 6AZ1 with its accompanying map EMD:7024. **b, c**, Two of the top GlideEM poses with generated from PDB:6AZ1 and EMD:7024. **d-f**, Top poses from traditional docking into PDB: 6AZ1, overlaid with the map EMD:7024.

### Using GemSpot with 3.0-4.5 Å resolution maps

The bulk of the cryoEM maps of liganded complexes in the PDB at the time of this publication fall into the range of 3.0-4.5 Å. This is unsurprising, as it is still challenging to achieve sub-3.0 Å EM maps, whereas maps worse than 4.5 Å are unlikely to present interpretable densities for the ligand. It is also within this resolution range that we expect modelers to encounter the greatest difficulty placing ligands by hand. Using GlideEM we were able to identify candidate poses in this resolution range with cross correlations that were comparable to the deposited poses for all of the structures that we studied. For GABA_A_ in complex with three different ligands^37^, the M2 muscarinic acetylcholine receptor in complex with two different ligands^38^, the sodium channel Nav1.7 with one ligand^39^, ATP citrate lyase with one ligand^7^, and the serotonin transporter with two different ligands^40^ the poses we identified with GlideEM largely agreed with what was deposited in the PDB (Supplementary Fig. 7) (although it should be noted that for the deposited structures of the M2 receptor and serotonin transporter the ligands were docked with the traditional version of Glide). At this resolution it is extremely unlikely to resolve water molecules. Accordingly, although our JAWS calculations predicted tightly bound water molecules in these structures, no such water molecules were located in the deposited maps.

In the example of the TRPM8 channel, the modeled pose for the ligand in 6NR3^41^ agrees well with what we predicted by GlideEM, although this represents another example where docking without the EM map yields additional poses that fall elsewhere in the relatively large and open binding site. In contrast, the icilin/TRPM8 model in 6NR2^41^ has significant strain energy in its ligand conformation. Using implicit solvent QM minimization we found that the deposited conformation for icilin is not a minimum energy conformation, but instead a much more linear conformation is energetically preferred (Fig. 4d). Comparison of the QM energy for the implicit solvent minimized ligand to that minimized with the strained dihedral fixed shows the restrained pose to be higher in energy by 5.2 kcal/mol. By employing GlideEM we were able to identify poses with significantly improved real-space cross correlations to the map (0.76-0.79, compared to 0.67 for the deposited pose) and without the heavily distorted dihedral (Fig. 4b,c).

**Fig 4.**
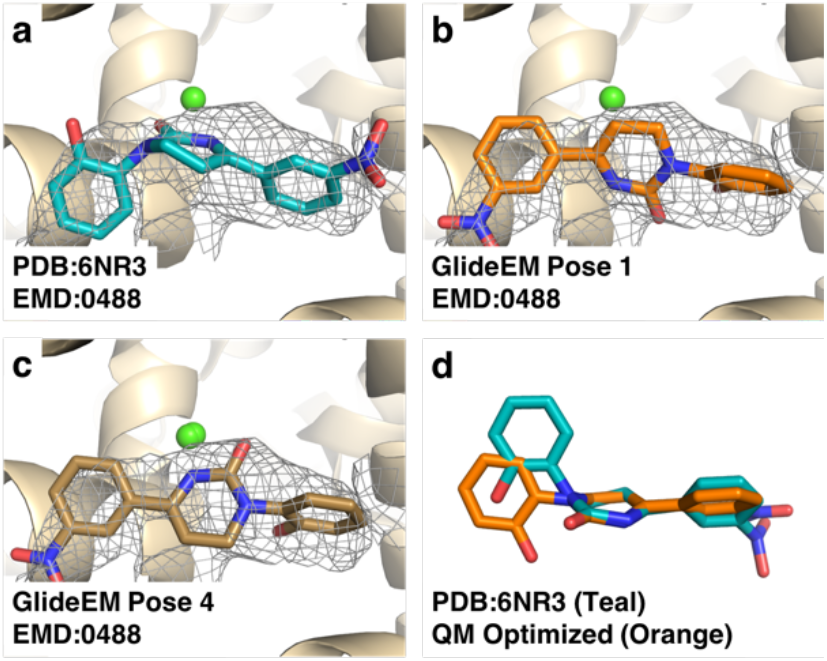
Comparison of poses for icilin bound to TRPM8. **a**, Deposited pose of icilin, PDB: 6NR3 with map EMD:0488. **b, c**, Two of the top docked GlideEM poses for icilin. **d**, The icilin conformation from the pose in 6NR3 in teal overlaid with its QM implicit solvent optimized pose in orange; both molecules are aligned to their central ring.

As the results with deposited structures in the 3.0-4.5 Å range were favorable, we wanted to explore how far we could push the low resolution end for the GemSpot pipeline. To this end we focused on the *leishmania* ribosome system, which as mentioned previously, does not find the correct ligand pose if the EM density is not used in docking. By taking a random subset of 15,000; 5,000; and 2,500 particles we calculated, respectively, 3.6. 4.3, and 5.5 Å resolution maps of the *leishmania* ribosome small subunit^35^. These lower resolution maps were then used to test the limits of the GlideEM method. With these maps, each pose was almost identical to its matched pose docked with the higher resolution map, with the exception of the most solvent exposed ring (Fig. 5), although at 5.5 Å some greater deviation in all rings was observed. It has been suggested previously that for methods of automatic protein model building with cryoEM maps, regions where there is greater divergence between automatically generated models may represent either regions of increased flexibility and/or greater uncertainty in the map, and this is most likely the case for our method^42^.

**Fig 5.**
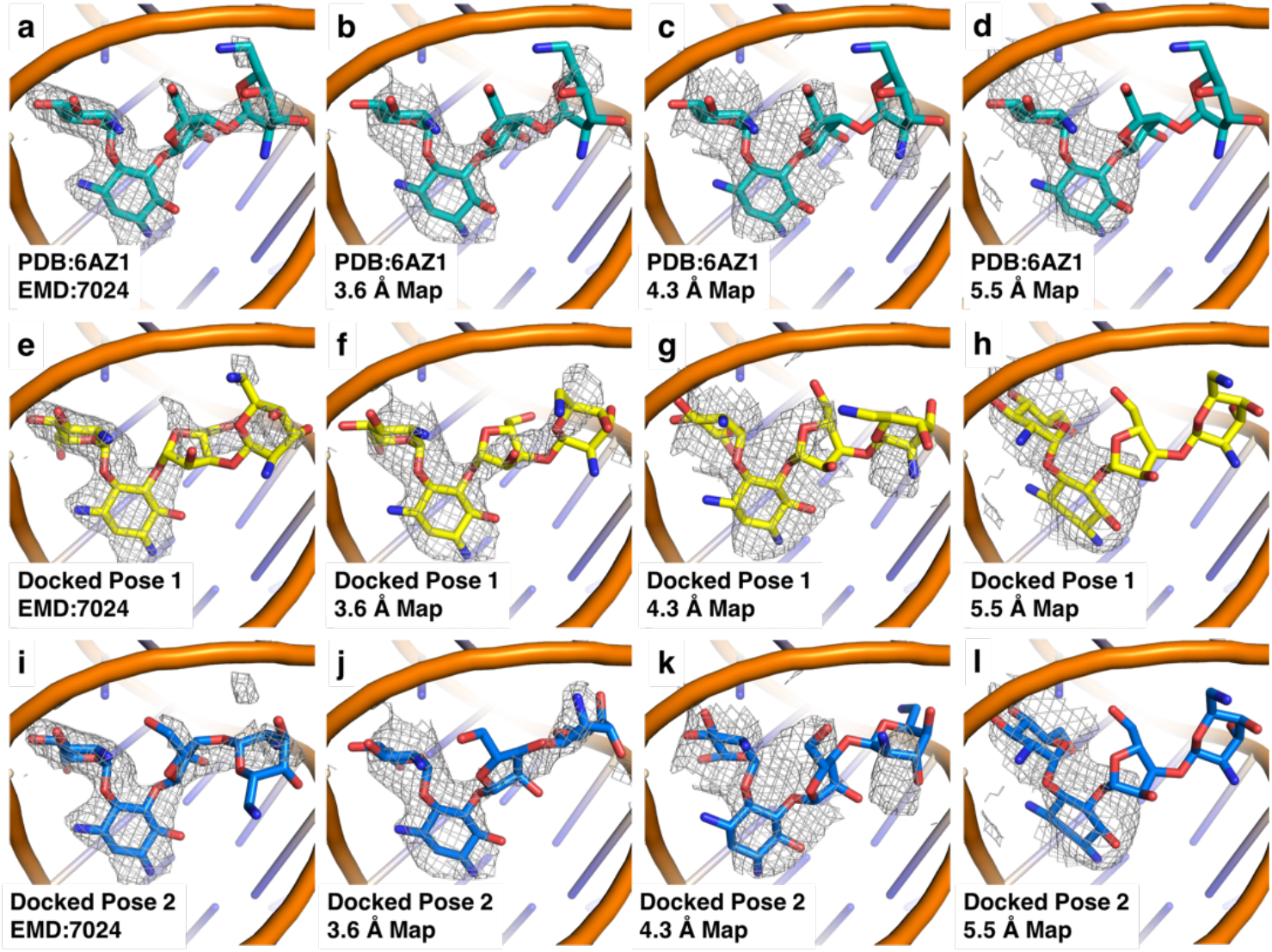
Comparison of poses for paromomycin bound to the *leishmania* ribosome in maps with global indicated resolutions of 2.6 Å, 3.6 Å, 4.3 Å, and 5.5 Å. a, b, c. Paromomycin pose from PDB: 6AZ1 with the maps at **a**, 2.6 Å **b**, 3.6 Å **c**, 4.3 Å **d**, 5.5 Å global resolution. **e, i**, The top two paromomycin poses from GlideEM using the original 2.6 Å map. **f, j**, The top two paromomycin poses from GlideEM using the 3.6 Å map from 15,000 particles. **g, k**, The top two paromomycin poses from GlideEM using the 4.3 Å map from 5,000 particles. **h, l**, The top two paromomycin poses from GlideEM using the 5.5 Å map from 2,500 particles.

### Using GemSpot with peptide ligands

With a thorough understanding of how the GemSpot pipeline performs for maps of various resolutions, we wanted to ensure the pipeline has good coverage of chemical space, particularly not just small molecule ligands but also peptides. Peptides often present a challenging class of ligands for computational docking because they have significantly more degrees of freedom than most druglike small molecules and thus require specialized protocols. Similarly, the standard implementation of GlideEM does not perform well on these molecules, and we thus developed a variant of the GlideEM based on the Glide peptide docking variant^43^ (details of the implementation are given in the methods). This methodology was applied to two liganded macromolecular complexes, that of the DAMGO (Tyr-D-Ala-Gly-NMePhe-Gly-ol) bound to the μ-opioid receptor/Gi complex^44^ and that of JMV449 (Lysψ(CH2NH)Lys-Pro-Tyr-Ile-Leu-OH) bound to the neurotensin type 1 receptor/Gi complex^45^ (Fig. 6). In both systems, we identified a pose nearly identical to the manually modeled one, as well as other poses that are consistent with the EM map as assessed by cross-correlation. This case necessitates the usage of additional information to choose an optimal pose. For example, SAR data across μ-opioid ligands suggests a highly conserved protonated amine group that interacts with aspartate 3.32^44^. DAMGO also has a protonated amine, and of the three poses with high real-space cross correlations only one has a salt bridge with aspartate 3.32 (Figure 6b), as is the case in the deposited pose (Figure 6a). Thus, we would suggest this to be the most probable experimental pose for DAMGO. While additional structures with larger peptides are necessary to more robustly validate GlideEM for peptides, this initial proof of concept provides a promising starting point.

**Fig 6.**
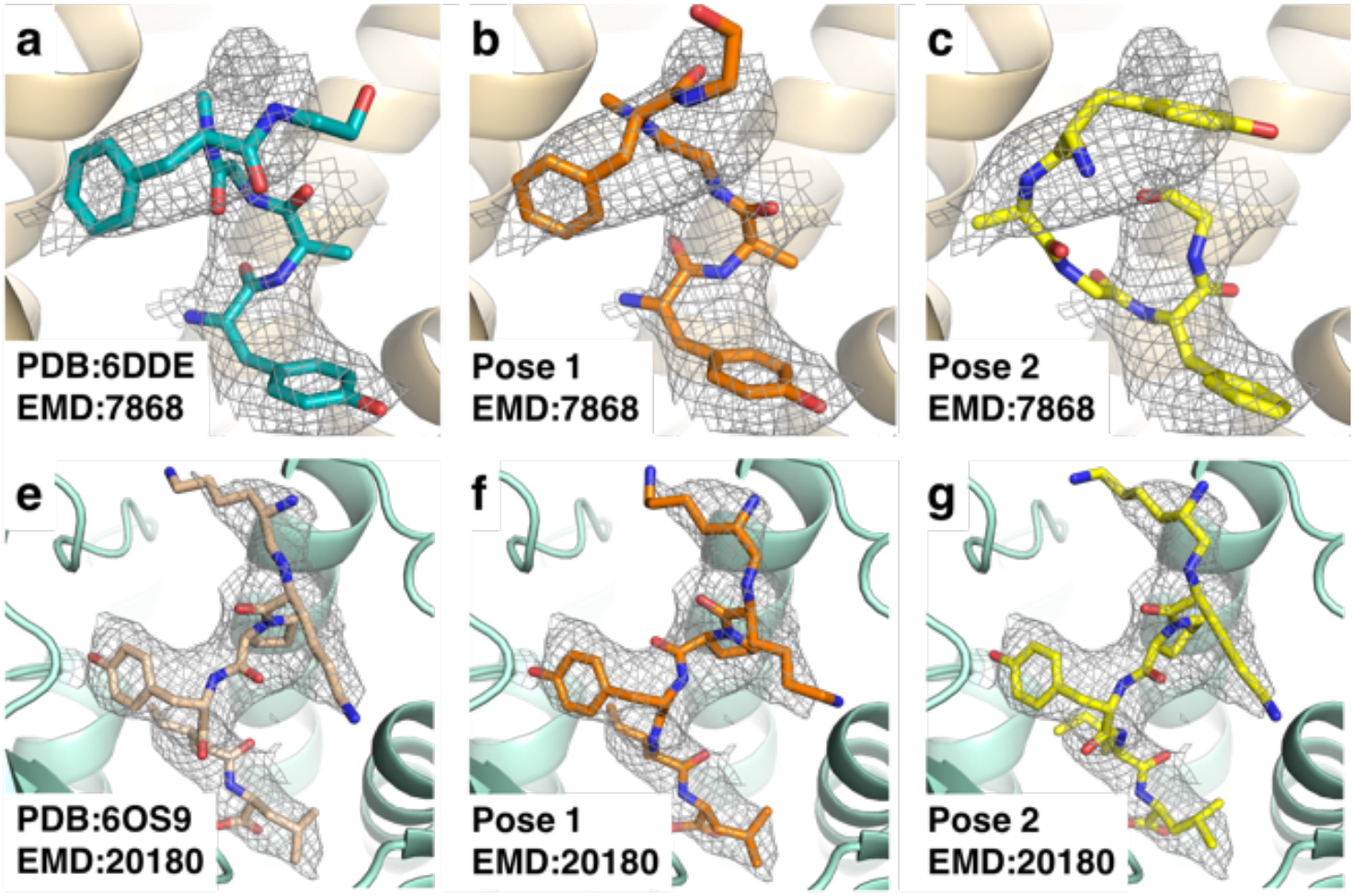
Results of docking peptides with GlideEM. **a**, Deposited structure of the μ-opioid-DAMGO complex, PDB:6DDE, EMD:7868. **b, c**, The two top poses for DAMGO from GlideEM. **d**, Deposited structure for the neurotensin type **1** receptor-JMV449 complex, PDB:6OS9, EMD:20180. **e, f**, The two top poses for JMV449 from GlideEM.

### Caveats and problematic cases

Although for most of ligand complex systems tested here the GemSpot pipeline works very well, some cases presented challenges that required special attention. While for the majority of structures the results were the same if the initial protein structure was refined with an empty ligand pocket or with some pose of the ligand present, in roughly 10% of systems an empty pocket would cause protein atoms to move into the map corresponding to the ligand, thereby preventing a correct pose from being properly docked. One example of this was GABA_A_ in complex with benzodiazepine ligands, where protein refinement with an empty ligand pocket lead to side chain atoms slipping into the ligand portion of the map during refinement (Supplementary Fig. 8). It is thus important that the modeler checks the structure and map before using GlideEM to ensure the protein is modeled optimally. This could also perhaps be remedied with an induced-fit docking approach where protein motions are informed by the EM map, although this is beyond the scope of the current study.

## Conclusion

As cryoEM has started to provide near-atomic and even atomic level detail of macromolecular complexes that proved impenetrable to traditional structural biology, it is becoming increasingly important to develop new tools ensuring that the most accurate structures are modeled into the experimental maps. Here we have presented and validated GemSpot, a workflow that combines computational chemistry methods with cryoEM maps to yield high-confidence models for the bound poses of ligands in macromolecular complexes. The novel GlideEM method will be made available in the newest version of the Schrödinger software package. We note, however, that the overall GemSpot approach described here should be implementable in any docking software package. We anticipate that GemSpot and its continuous evolvement will become an invaluable tool for the correct interpretation and modeling of ligand densities and will greatly aid in drug discovery efforts that are based on cryoEM.

## Online Methods

The GemSpot pipeline starts with preparing and refining structures and ligands, by loading each structure and ligand into Maestro where they were processed using the Protein Preparation Wizard panel^46^ with default options, i.e. missing side chain atoms and hydrogens were added, and the hydrogen bonding network optimized. For peptides an additional sampling step was performed to create a diverse set of backbone conformations using the Peptide Docking panel in Maestro, outputting 1000 conformations^43^. Next, the ligands were docked with GlideEM, a version of Glide that takes into account the cryoEM potential map.

Glide, similar to other docking algorithms^47,48^, starts by sampling the isolated ligand to determine an ensemble of low-energy conformations that could be biologically relevant. Each pose is rapidly scored on a grid with rewards for creating favorable protein-ligand interactions, such as hydrogen bonds and lipophilic interactions, and penalties for unfavorable ones, like steric clashes. Poses that score highly at this step are then refined by a combination of local sampling and minimization in a force field that includes ligand strain energy, as calculated by the force field, as well as protein-ligand interactions. For GlideEM we have added a simple real-space cross-correlation score used in both the sampling and refinement stages in order to help ensure that the docking function finds the correct binding mode given the experimental cryoEM data as follows.

The input CryoEM map is first normalized in the region around the binding site, by subtracting the mean and dividing by the standard deviation of intensities found over all voxels within a distance of 10 Ångstroms plus the ligand’s radius of gyration, determined during the conformational sampling stage, from the center of the binding site. The normalized map is used in the sampling step by rewarding the placement of non-hydrogen atoms in areas with higher Coulombic potential in addition to its standard scoring function. During the refinement step, the protein-ligand energy function is augmented by the real space cross-correlation between the normalized map and a simple simulated potential map of the ligand where each atom is modeled as a single Gaussian function for computational efficiency. Each pose is ultimately ranked with a combination of the standard Glide energy function, the *GlideScore*, and the approximate fit to the experimental density, the *DensScore*, to determine the poses that will be evaluated with a more rigorous scoring function. Two parameters are exposed to the user to control the weight of scoring against the EM map, *faceden* and *facrf*. The *facrf* parameter controls the weight of the EM map during the initial sampling phase, and *faceden* during the refinement phase. While high values tend to increase the contribution of the EM map too much and generally lead to highly strained conformations of the ligand, at present there is an insufficient number of deposited structures to rigorously determine optimal parameters. Thus, both *facrf* and *faceden* were set to 1 for all systems studied in this work. As with traditional Glide, an Emodel is also provided for each pose. This value is the result of a scoring function designed to better rank different poses of the same compound, and may be useful in cases where Glide score and cross-correlation alone provide more than one equally ranked pose.

The next step in the pipeline consists of refining top scoring poses generated by GlideEM. For small molecules the top 5 poses with the best GlideScore were chosen. For our peptide protocol, poses were picked based on the DensScore, starting with the best scoring pose and adding additional poses that have a heavy atom RMSD higher than 0.5 Å compared to already picked poses up to a total set of 100. The chosen poses were refined with PHENIX’s real space refinement using an adjusted protocol where the OPLS3e /VSGB2.1 force field is used to calculate ligand energies. Here the macromolecule was processed separately from the ligand of interest with phenix.ready_set (default parameters), and recombined with the ligand poses coming from GlideEM. CIF files for the ligands were created using the hetgrp_ffgen Schrödinger utility. Although the parameters of the ligands’ CIF files are unimportant for the energy model, they are essential for PHENIX to run correctly, and needed to provide the ligand’s topology for restraint B-factor refinement. During real space refinement with phenix.real_space_refine, the chemical energy components of the ligands were swapped with the OPLS3e / VSGB2.1 force field energies with a weight factor of 10 to place the forces on the same order of magnitude of the default PHENIX restraints model. Otherwise, default parameters were used during refinement. All real-space cross correlations were calculated with phenix.cryoem_validation.

The final steps in the GemSpot pipeline, after docking and refinement, consist of further analyzing the generated ligand conformations using quantum chemical calculations, water placement simulations, and including known SAR if available. Quantum chemical calculations were performed with the GAUSSIAN software^28^. Ligand structures were minimized with the ωB97-xd functional^49^ and the 6-311+(2d,2p) basis set in SMD implicit water, optimizing all degrees of freedom.

JAWS Monte Carlo simulations^22^ were set up as follows. The refined structures were culled at a 25 Å sphere centered around the ligand’s center of mass. The system was simulated in MCPRO^50^ with the OPLS-AA/M^51^ force field for the protein and OPLS-AA/CM1A for the ligands. The ligand was solvated with a 5 Å layer of TIP4P^52^ theta-water and a 25Å spherical cap of TIP4P water beyond that. Sidechains within 15Å of the ligand were allowed to sample flexibly. 5 million Monte Carlo steps were used for solvent equilibration, 10 million in hydration site identification, and 50 million for production. Three independent simulations were run for each system with the strong consensus water molecules (predicted binding affinity better than 3 kcal/mol) used to locate average positions for subsequent PHENIX real space refinement using the protocol outlined above.

For this work, the systems chosen were phenethyl beta-d-thiogalactoside (PETG)/beta-galactosidase (PDB:5A1A, EMD:2984)^30^; PETG/beta-galactosidase (PDB:6CVM, EMD:7770)^14^; Tetrodotoxin/Voltage gated sodium channel NavPaS (PDB:6A95, EMD:6995)^34^; Fubinaca/Cannabinoid receptor 1 (PDB:6N4B, EMD:0339)^33^; GSK 3494245*/Leishmania* 20S proteasome (PDB:6QM7, EMD:4590)^36^; Paromomycin/*Leishmania* ribosome (PDB:6AZ1, EMD:7024)^35^; Menthol analogue WS-12/Ion channel TRPM8 (PDB:6NR2, EMD:0488)^41^; Icilin/Ion channel TRPM8 (PDB:6NR3, EMD:0487)^41^; LY2119620/M2R (PDB:6OIK, EMD:20079)^38^; Iperoxo/M2R (PDB:6OIK, EMD:20079)^38^; Saxitoxin/Nav1.7 (PDB: 6J8G, EMD:9781)^39^; Biculine/GABAA (PDB:6HUK, EMD:0280)^37^; Xanax/GABAA (PDB:6HUO, EMD:0282)^37^; Valium, GABA_A_ (PDB:6HUP, EMD:02 83)^37^; NDI-091143/ATP Citrate Lyase (PDB:6O0H, EMD:0567)^7^; paroxetine/serotonin transporter (PDB:6DZW, EMD:8941)^40^; ibogaine/serotonin transporter (PDB:6DZZ, EMD:8943)^40^; ibogaine, serotonin transporter (PDB:6DZY,EMD:8942)^40^; DAMGO/Mu opioid receptor (PDB:6DDE, EMD:7868)^44^, and JMV449/Neurotensin receptor (PDB:6OS9, EMD:20180)^45^.

## Software

The following software packages were used: Maestro and the Schrödinger software stack (a modified build of the 2019-3 distribution; https://www.schrodinger.com/downloads/releases) for prepping and docking compounds. PHENIX (v1.15; http://phenix-online.org/download/) for real space refinement (in combination with Schrödinger for the OPLS3e / VSGB2.1 force field) and cross correlation calculations. GAUSSIAN (v16) for quantum chemical energy calculations; and MCPRO (2.3) for hydrating and detecting water molecules; PyMol (2.3.0) was used for generating figures.

## Supporting information

Full SI

## Acknowledgments

We would like to thank William Weis and Axel Brunger for comments on the manuscript. Author Contributions M. J. R and G. S. initiated the project. G. C. P. v. Z. and K. B. and developed & implemented software. M. J. R. and K. B. ran docking and refinement calculations. M. J. R. performed QM and JAWS calculations. M. J. R. and G. S., wrote the manuscript with input from K. B. and G. C. P. v. Z.

