## Supplementary material for "GemSpot: A Pipeline for Robust Modeling of Ligands into CryoEM Maps": Full SI

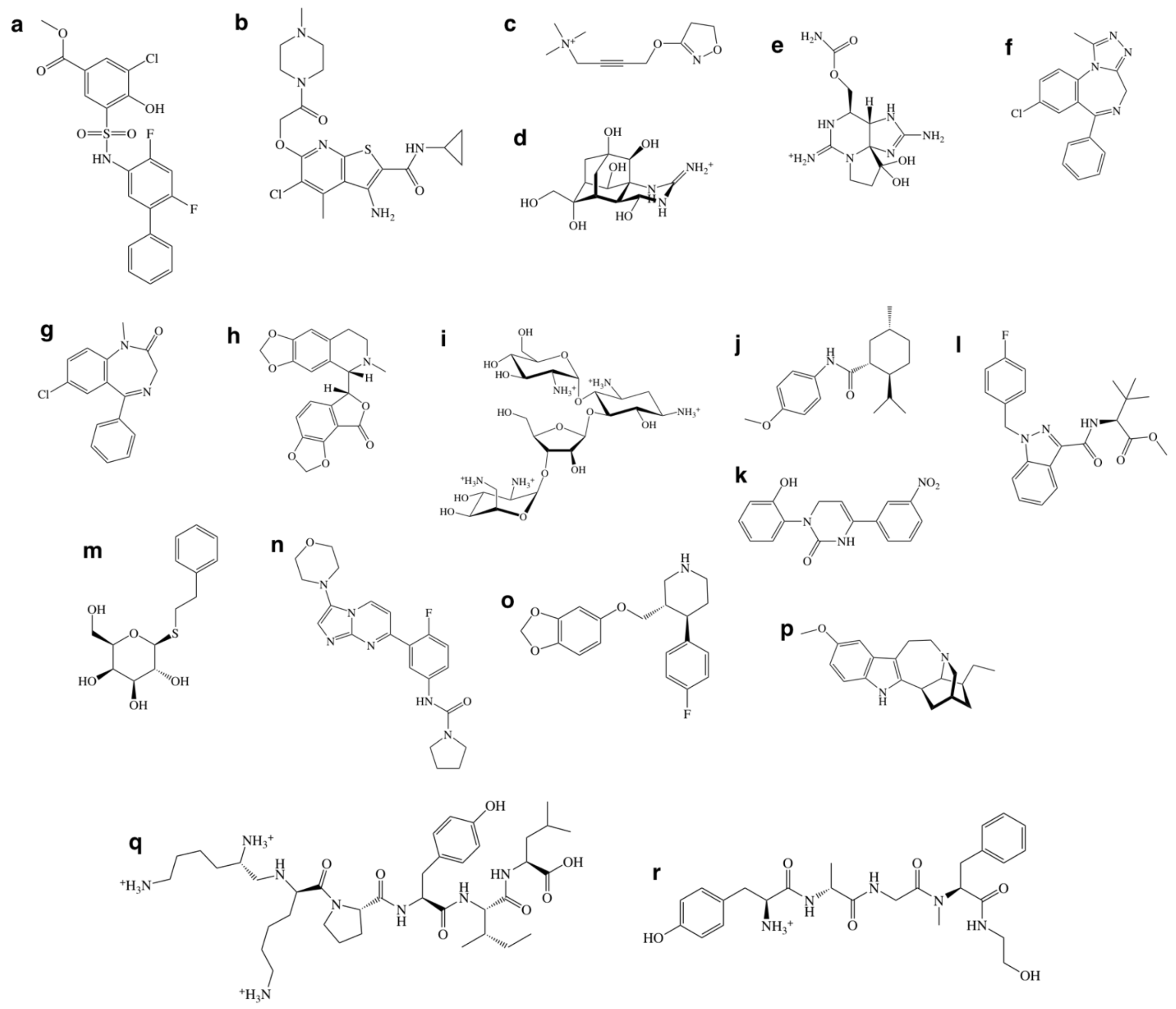

**Supplementary Figure 1: Ligands docked in this work.**

**a,** NDI-091143/ATP citrate lyase. **b,** LY2119620/M2 receptor. **c,** Iperoxo/M2 receptor. **d,** Tetrodotoxin/NaVaPas. **e,** Saxitoxin/NaV1.7. **f,** Xanax/$GABA_A$. **g,** Valium/$GABA_A$. **h,** Biculine/$GABA_A$. **i,** Paromomycin/*leishmania* ribosome. **j,** Menthol analogue WS-12/TRPM8. **k,** Icilin/TRPM8. **l,** Fubinaca/cannabinoid type 1 receptor. **m,** PETG/beta-galactosidase. **n,** GSK3494245/*leishmania* 20S proteasome. **o,** Paroxetine/serotonin transporter. **p,** Ibogaine/serotonin transporter. **q,** DAMGO/μ-opioid receptor. **r,** JMV449/neurotensin type 1 receptor.

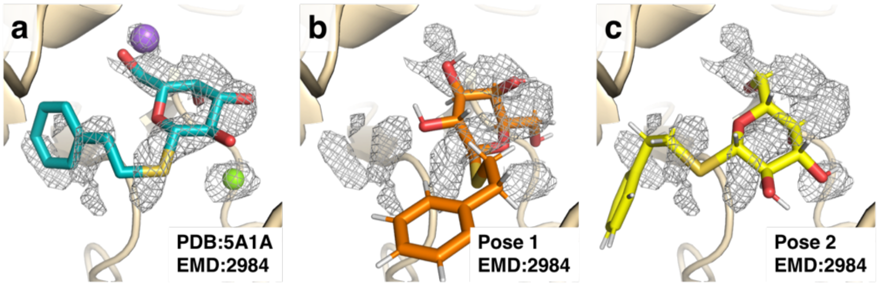

**Supplementary Figure 2**: **PETG poses docked without EM map densities.**

**a,** Deposited structure PDB:5A1A PETG bound to beta-galactosidase, with displayed density from EMD:2984. The purple and green sphere correspond to a sodium and magnesium ion, respectively. **b,** Top pose for PETG from docking to PDB:5A1A without using the EM density, overlaid with EMD:2984. **c,** Another pose from docking to PDB:5A1A without density, overlaid with EMD:2984.

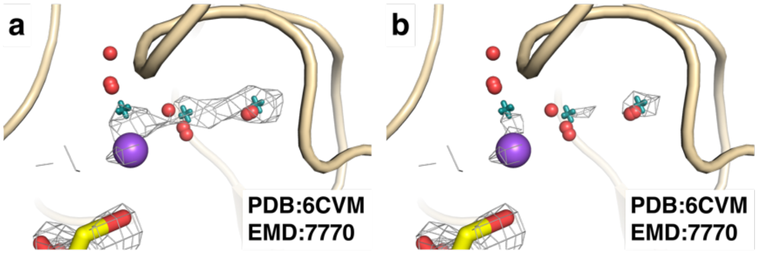

**Supplementary Figure 3**: **Additional beta-galactosidase hydration sites predicted by JAWS.**

Predicted sites of hydration from triplicate JAWS simulations on NavPaS are represented as red spheres, while locations of real-space refined waters are represented as teal crosses. The purple sphere corresponds to a sodium ion. **a, b,** Show density for EMD:7770 at two different contour levels.

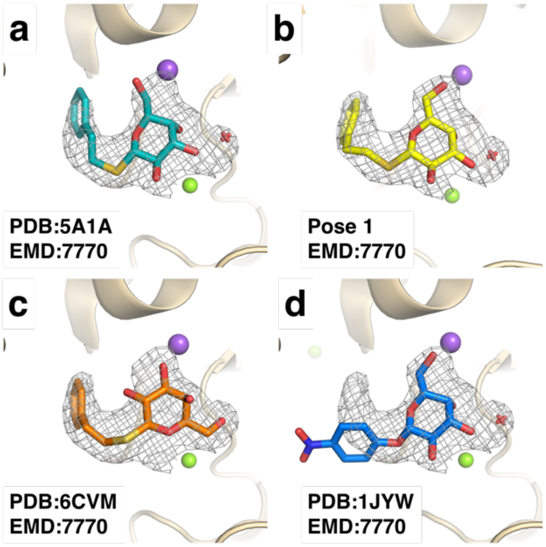

**Supplementary Figure 4: Comparison of the PETG poses from cryoEM and a high-resolution crystal structure of an analogue.**

All panels display map density from EMD:7770. Purple spheres, green spheres, and red crosses correspond to sodium ions, magnesium ions, and water molecules, respectively. **a,** PDB: 5A1A. **b,** Our best GemSpot pose with PDB: 6CVM and EMD:7770. **c,** PDB: 6CVM. **d,** The high-resolution crystal structure of analogous compound PNPG (PDB:1JYW).

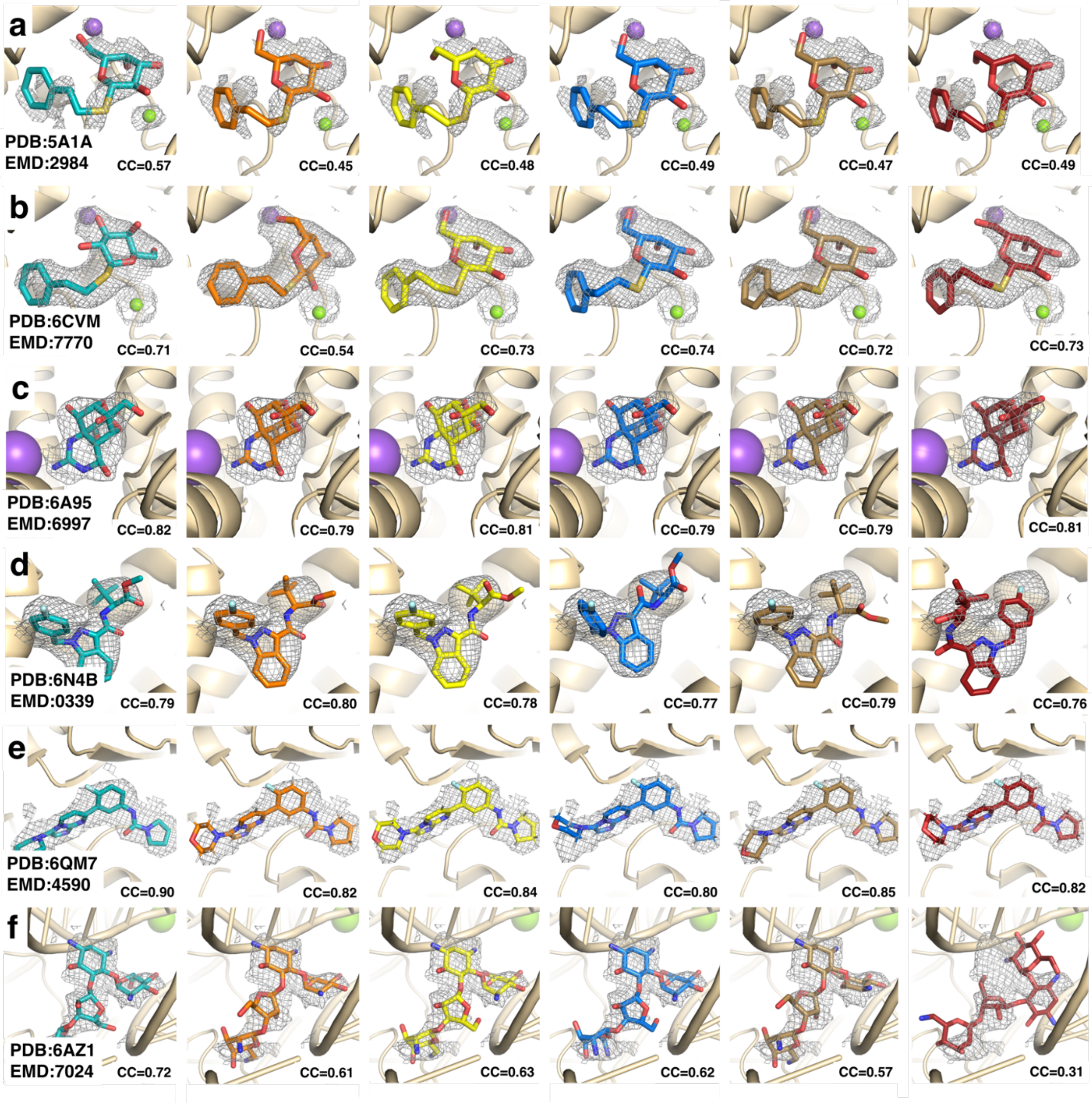

**Supplementary Figure 5**: **Sub-3.0 Å cryoEM maps from this work**.

In all cases the deposited pose in the PDB is in teal while the orange, yellow, blue, brown, and red poses in the same row correspond to pose 1-5, respectively, in supplementary table 2. CC is the real space cross-correlation calculated without hydrogen atoms. **a,** PETG/beta-galactosidase (PDB:5A1A, EMD:2984). **b,** PETG/beta-galactosidase (PDB:6CVM, EMD:7770). **c,** Tetrodotoxin/NavPaS (PDB:6A95, EMD:6995). **d,** Fubinaca/cannabinoid type 1 receptor (PDB:6N4B, EMD:0339). **e,** GSK 3494245/*leishmania* 20S proteasome (PDB:6QM7, EMD:4590). **f,** Paromomycin/*leishmania* ribosome (PDB:6AZ1, EMD:7024).

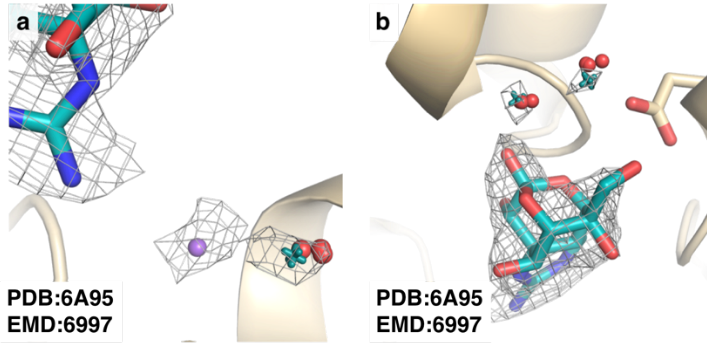

**Supplementary Figure 6: Predicted water sites in the structure of tetrodotoxin bound to NavPaS.**

The EM density map in wireframe corresponds to that of EMD:6995. Red spheres represent predicted sites of hydration from triplicate JAWS runs while teal crosses represent water molecule locations after real space refinement. **a,** The sodium ion (purple sphere) and its bound water molecule. **b,** Two additional probable sites of hydration.

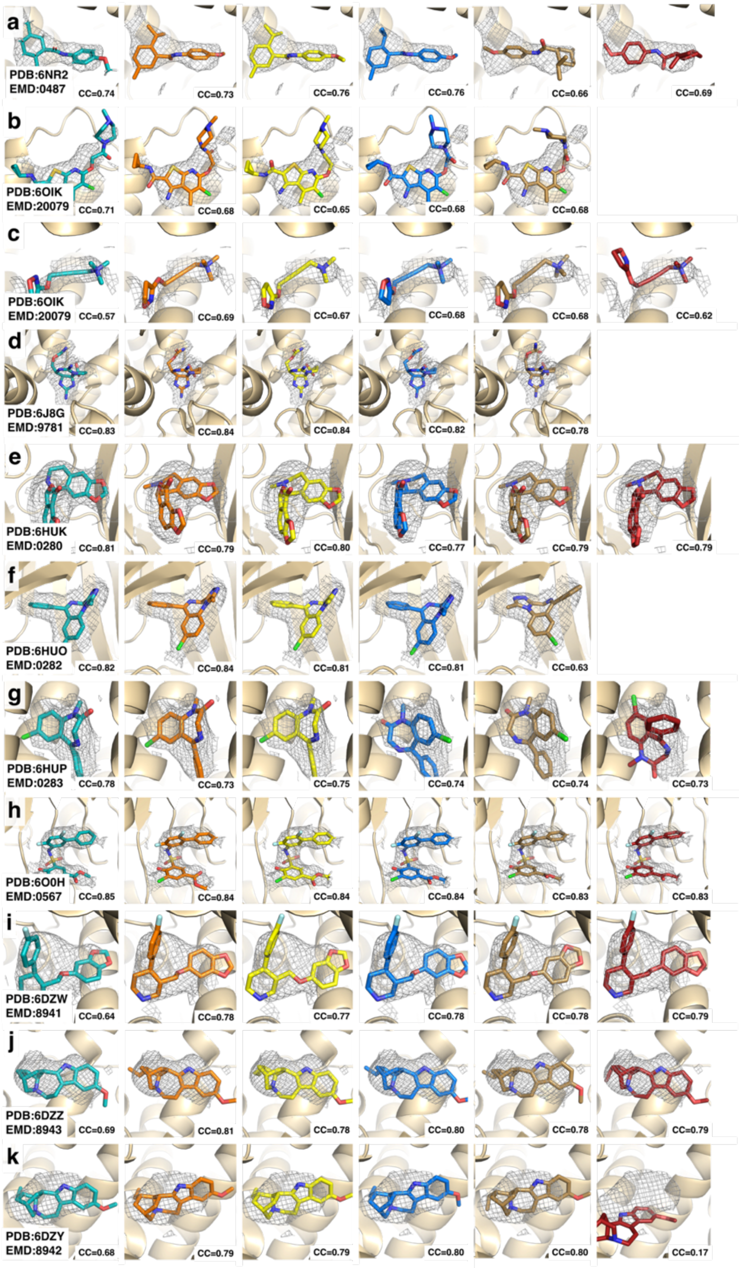

**Supplementary Figure 7**: **3.0 Å - 4.5 Å maps from this work**.

In all cases the deposited pose in the PDB is in teal while the orange, yellow, blue, brown, and red poses in the same row correspond to pose 1-5, respectively, in supplementary table 2. CC is the real space cross-correlation calculated without hydrogen atoms. **a,** Menthol analogue WS-12/TRPM8 (PDB:6NR2, EMD:0487). **b,** LY2119620/M2R (PDB:6OIK, EMD:20079). **c,** Iperoxo/M2R (PDB:6OIK, EMD:20079). **d,** Saxitoxin/Nav1.7 (PDB: 6J8G, EMD:9781). **e,** Biculine/$GABA_A$ (PDB:6HUK, EMD:0280). **f,** Xanax/$GABA_A$ (PDB:6HUO, EMD:0282). **g,** Valium, $GABA_A$ (PDB:6HUP, EMD:0283). **h,** NDI-091143/ATP Citrate Lyase (PDB:6O0H, EMD:0567). **i,** Paroxetine/serotonin transporter (PDB:6DZW, EMD:8941). **j,** Ibogaine/serotonin transporter (PDB:6DZZ, EMD:8943). **k,** Ibogaine, serotonin transporter (PDB:6DZ, EMD:8942).

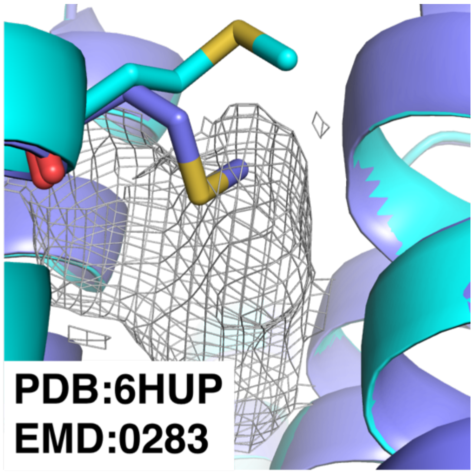

**Supplementary Figure 8: Comparison of PDB:6HUP refined with and without ligand present.**

$GABA_A$ receptor structure PDB:6HUP shown in cyan; the re-refined structure in the absence of ligand is shown in blue. The map corresponds to the portion of EMD:0283 corresponding to the bound ligand.

**Supplementary Table 1**

All systems studied in this work, sorted by resolution.

| Name | PDB Code | Resolution | Biomolecule | Ligand |
|---|---|---|---|---|
| Beta-Galactosidase | 6CVM | 1.9 | Protein | Phenethyl β-d-thiogalactoside |
| Beta-Galactosidase | 5A1A | 2.2 | Protein | Phenethyl β-d-thiogalactoside |
| NavPaS | 6A95 | 2.6 | Membrane Protein | Tetrodotoxin |
| Leishmania Ribosome Small Subunit | 6AZ1 | 2.7 | RNA/Protein | Paromomycin |
| Leishmania 20S Proteosome | 6QM7 | 2.8 | Protein | GSK3494245 |
| Cannabinoid Receptor 1 | 6N4B | 3.0 | Membrane Protein | Fubinaca |
| Nav1.7 | 6J8G | 3.2 | Membrane Protein | Saxitoxin |
| GABA$_A$ Receptor | 6HUO | 3.26 | Membrane Protein | Xanax |
| TRPM8 | 6NR3 | 3.4 | Membrane Protein | Icilin |
| Mu Opioid Receptor | 6DDF | 3.5 | Membrane Protein | DAMGO |
| GABA$_A$ Receptor | 6HUP | 3.58 | Membrane Protein | Valium |
| Muscarinic Receptor 2 | 6OIK | 3.6 | Membrane Protein | Iperoxo |
| Muscarinic Receptor 2 | 6OIK | 3.6 | Membrane Protein | LY2119620 |
| Serotonin Transporter | 6DZZ | 3.6 | Membrane Protein | Ibogaine |
| ATP Citrate Lyase | 6O0H | 3.67 | Protein | NDI-091143 |
| GABA$_A$ Receptor | 6HUK | 3.69 | Membrane Protein | Biculine |
| TRPM8 | 6NR2 | 4.0 | Membrane Protein | Menthol Analogue WS-12 |
| Serotonin Transporter | 6DZY | 4.1 | Membrane Protein | Ibogaine |
| Serotonin Transporter | 6DZW | 4.3 | Membrane Protein | Paroxetine |

### Supplementary Table 2

Docking and refinement results for the systems studied in this work.

| System | Pose | Glide | Emodel | CC Post Refinement | CC Post Refinement (non-hydrogen atoms) | Deposited Pose CC |
|---|---|---|---|---|---|---|
| Bgal | 1 | -5.6 | -136.7 | 0.49 | 0.45 | 0.57 |
| PETG | 2 | -5.4 | -137.9 | 0.51 | 0.48 | |
| 5A1A | 3 | -5.3 | -132.9 | 0.52 | 0.49 | |
| | 4 | -5.3 | -133.7 | 0.51 | 0.47 | |
| | 5 | -4.7 | -134.4 | 0.51 | 0.49 | |
| Bgal | 1 | -9.9 | -240.9 | 0.63 | 0.54 | 0.71 |
| PETG | 2 | -7.9 | -328.3 | 0.74 | 0.73 | |
| 6CVM | 3 | -7.4 | -315.6 | 0.75 | 0.74 | |
| | 4 | -7.4 | -309.3 | 0.73 | 0.72 | |
| | 5 | -5.6 | -308.2 | 0.74 | 0.73 | |
| NavPaS | 1 | -5 | -1226.9 | 0.75 | 0.79 | 0.82 |
| Tetrodotoxin | 2 | -4.9 | -1217.1 | 0.77 | 0.81 | |
| 6A95 | 3 | -4.8 | -1226.7 | 0.73 | 0.79 | |
| | 4 | -4.6 | -1226.9 | 0.73 | 0.79 | |
| | 5 | -4.4 | -1218.9 | 0.68 | 0.81 | |
| TRPM8 | 1 | -5 | -239.3 | 0.73 | 0.73 | 0.74 |
| Menthol Analogue WS-12 | 2 | -4.9 | -239.2 | 0.74 | 0.76 | |
| 6NR2 | 3 | -4.3 | -244.3 | 0.74 | 0.76 | |
| | 4 | -3.8 | -249.3 | 0.69 | 0.66 | |
| | 5 | -3.8 | -240.2 | 0.69 | 0.69 | |
| TRPM8 | 1 | -6.1 | -303.5 | 0.77 | 0.79 | 0.67 |
| Icilin | 2 | -6 | -311.7 | 0.75 | 0.78 | |
| 6NR3 | 3 | -6 | -319.3 | 0.75 | 0.77 | |
| | 4 | -5.6 | -303.5 | 0.8 | 0.79 | |
| | 5 | -4.6 | -305.1 | 0.75 | 0.76 | |
| Cannabinoid Receptor 1 | 1 | -9.4 | -474.3 | 0.74 | 0.80 | 0.79 |
| Fubinaca | 2 | -9.1 | -464.3 | 0.74 | 0.78 | |
| 6N4B | 3 | -8.9 | -464.6 | 0.73 | 0.75 | |
| | 4 | -8.8 | -475.4 | 0.72 | 0.73 | |
| | 5 | 18.7 | 3325.6 | 0.52 | 0.49 | |
| GABA(A) | 1 | -7.9 | -839 | 0.82 | 0.84 | 0.82 |
| Xanax | 2 | -7.6 | -803 | 0.79 | 0.81 | |
| 6HUO | 3 | -7.3 | -815 | 0.79 | 0.81 | |
| | 4 | -5.4 | -774 | 0.62 | 0.63 | |
| GABA(A) | 1 | -8.4 | -842 | 0.78 | 0.79 | 0.81 |
| Biculine | 2 | -8.1 | -876 | 0.79 | 0.80 | |
| 6HUK | 3 | -7.8 | -807 | 0.78 | 0.77 | |
| | 4 | -7.8 | -879 | 0.77 | 0.79 | |
| | 5 | -7.5 | -800 | 0.78 | 0.79 | |
| GABA(A) | 1 | -7.9 | -408.2 | 0.75 | 0.73 | 0.78 |
| Valium | 2 | -6.7 | -434.1 | 0.75 | 0.75 | |
| 6HUP | 3 | -6 | -418.3 | 0.75 | 0.74 | |
| | 4 | -5.3 | -402.4 | 0.73 | 0.74 | |
| | 5 | -5.1 | -413.3 | 0.75 | 0.73 | |
| Muscarinic Receptor 2 | 1 | -8.3 | -687.3 | 0.66 | 0.68 | 0.71 |
| LY2119620 | 2 | -7.4 | -687.3 | 0.63 | 0.65 | |

| | | | | | | |
|---|---|---|---|---|---|---|
| 6OIK | 3 | -7.3 | -694.8 | 0.64 | 0.68 | |
| | 4 | -6.6 | -678.5 | 0.64 | 0.68 | |
| Muscarinic Receptor 2 | 1 | -8.3 | -161.6 | 0.73 | 0.69 | 0.57 |
| Iperoxo | 2 | -7.7 | -165.8 | 0.71 | 0.67 | |
| 6OIK | 3 | -7.3 | -159 | 0.72 | 0.68 | |
| | 4 | -7.2 | -159.7 | 0.7 | 0.68 | |
| | 5 | -6.8 | -111.2 | 0.68 | 0.62 | |
| ATP-Citrate Lyase | 1 | -9.3 | -665.7 | 0.85 | 0.84 | 0.85 |
| NDI-091143 | 2 | -8.8 | -643.4 | 0.84 | 0.84 | |
| 6O0H | 3 | -8.7 | -680.7 | 0.84 | 0.84 | |
| | 4 | -7.5 | -681.6 | 0.84 | 0.83 | |
| | 5 | -7.2 | -652.2 | 0.85 | 0.83 | |
| Nav1.7 | 1 | -5.5 | -627.8 | 0.81 | 0.84 | 0.83 |
| Saxitoxin | 2 | -5.4 | -692.9 | 0.82 | 0.84 | |
| 6J8G | 3 | -5 | -555.9 | 0.79 | 0.82 | |
| | 4 | -3.2 | -608 | 0.77 | 0.78 | |
| Leishmania 20S Proteasome | 1 | -6.6 | -715.1 | 0.79 | 0.82 | 0.90 |
| GSK3494245 | 2 | -6.5 | -799.5 | 0.82 | 0.84 | |
| 6QM7 | 3 | -6.4 | -704.7 | 0.76 | 0.80 | |
| | 4 | -6.3 | -769.2 | 0.8 | 0.85 | |
| | 5 | -5.8 | -714.5 | 0.78 | 0.82 | |
| Serotonin Transporter | 1 | -7.5 | -393.8 | 0.79 | 0.78 | 0.64 |
| Paroxetine | 2 | -7.3 | -378 | 0.78 | 0.77 | |
| 6DZW | 3 | -6.7 | -371.4 | 0.79 | 0.78 | |
| | 4 | -6.2 | -388.9 | 0.8 | 0.78 | |
| | 5 | -4.2 | -379.4 | 0.8 | 0.79 | |
| Serotonin Transporter | 1 | -8 | -452.8 | 0.77 | 0.79 | 0.68 |
| Ibogaine | 2 | -7.4 | -417.8 | 0.77 | 0.79 | |
| 6DZY | 3 | -7.3 | -432.1 | 0.78 | 0.80 | |
| | 4 | -7 | -422 | 0.79 | 0.80 | |
| | 5 | -6.8 | -60.5 | 0.53 | 0.17 | |
| Serotonin Transporter | 1 | -7.4 | -534.7 | 0.81 | 0.81 | 0.69 |
| Ibogaine | 2 | -7.4 | -527.4 | 0.77 | 0.78 | |
| 6DZZ | 3 | -6.4 | -522.4 | 0.8 | 0.80 | |
| | 4 | -6.1 | -529.2 | 0.79 | 0.78 | |
| | 5 | -6 | -524 | 0.78 | 0.79 | |
| Leishmania Ribosome | 1 | -5.8 | -40645 | 0.70 | 0.61 | 0.72 |
| Paromomycin | 2 | -5.4 | -40687 | 0.68 | 0.63 | |
| 6AZ1 | 3 | -5.4 | -41242 | 0.65 | 0.62 | |
| | 4 | -4.7 | -41279 | 0.66 | 0.57 | |
| | 5 | -4.3 | -41017 | 0.39 | 0.31 | |
| Leishmania Ribosome | 1 | -5.8 | -52301 | 0.74 | | |
| Paromomycin | 2 | -5.5 | -51518 | 0.77 | | |
| 15k particles | 3 | -4.2 | -52797 | 0.76 | | |
| | 4 | -0.8 | -52075 | 0.77 | | |
| | 5 | 0.9 | -51741 | 0.76 | | |
| Leishmania Ribosome | 1 | -6.4 | -52255 | 0.80 | | |
| Paromomycin | 2 | -5.7 | -51587 | 0.79 | | |
| 5k particles | 3 | -5.6 | -51189 | 0.81 | | |
| | 4 | -5.7 | -51376 | 0.80 | | |
| | 5 | -1.3 | -52117 | 0.82 | | |
| Leishmania Ribosome | 1 | -5.6 | -43357 | 0.89 | | |
| Paromomycin | 2 | -4.5 | -46328 | 0.89 | | |
| 2.5k particles | 3 | -2.4 | -46599 | 0.89 | | |
| | 4 | 2.2 | -43985 | 0.89 | | |

| | | | | |
|---|---|---|---|---|
| | 5 | 2.9 | -43743 | 0.89 |
| Mu Opioid Receptor | | | | |
| DAMGO | 1 | -5.5 | -73.1 | 0.67 |
| 6DDE | 2 | -5.5 | -84.3 | 0.66 |
| | 3 | -5.7 | -95.2 | 0.63 |
| | 4 | -5.7 | -88.4 | 0.61 |
| | 5 | -5.5 | -74.8 | 0.59 |
| | 6 | -5.5 | -69.9 | 0.58 |
| | 7 | -5.4 | -82.4 | 0.57 |
| | 8 | -5.5 | -82.4 | 0.55 |

Glide and Emodel correspond to the Glide score and Emodel described in the methods section. CC is the cross correlation between the ligand model and the map.
